# Quantification of SARM1 activity in human peripheral blood mononuclear cells

**DOI:** 10.1101/2025.02.17.638666

**Authors:** Lila Dabill, Ivana Shen, Jennifer Brazill, Alicia Neiner, Yo Sasaki, Erica L Scheller

## Abstract

SARM1 (sterile α and TIR motif-containing protein-1) is an NADase enzyme that has been identified as the central executioner of Wallerian axon degeneration. Given this, SARM1 is of high interest as a candidate therapeutic target and SARM1 inhibitors are currently in clinical trials for prevention and treatment of neurodegeneration. Beyond neuroscience, emerging studies reveal that SARM1 activation may also drive aspects of bone fragility, liver pathology, adipose tissue expansion, and insulin resistance in settings of metabolic disease. However, we lack methods to quantify SARM1 activation in humans using clinical isolates to better define patients at high risk of SARM1-mediated tissue damage, informing the future clinical application of SARM1 inhibitors. Unlike neurons, peripheral blood mononuclear cells (PBMCs) represent an easily accessible population of cells for clinical screening. We hypothesized that by pairing activators and inhibitors of SARM1 with analysis of downstream changes in cellular metabolites, we could quantify both the basal SARM1 activity and the SARM1 activation potential of human PBMCs. Our results reveal that SARM1 agonist pyrinuron, also known as Vacor, activates a dose-dependent increase in cAPDR and the cADPR:ADPR ratio that is arrested when paired with SARM1 inhibitor DSRM-3716. Various changes in secondary metabolites were also characterized and reported herein. Overall, these findings demonstrate that human PBMCs have detectable SARM1 activation potential and could be leveraged as a clinical readout of SARM1 expression and activity across diverse disease contexts.

## INTRODUCTION

SARM1 (sterile α and TIR motif-containing protein-1) is an inducible NADase enzyme that is highly expressed in the nervous system. SARM1 serves as a metabolic biosensor and is activated in response to injury, inflammation, and oxidative stress.^1–4^ Based on a screen of thousands of candidates, SARM1 was identified as the central executioner of Wallerian axon degeneration.^4,5^ Since this time, SARM1 has also been shown to modulate nerve function through regulation of MAPK signaling and metabolite turnover.^6,7^ In response to this pivotal discovery, SARM1 inhibitors are now being developed for clinical management of neurodegenerative diseases.^8–10^ Beyond neuroscience, emerging studies reveal that SARM1 activation may also drive aspects of bone fragility, liver pathology, adipose tissue expansion, and insulin resistance in settings of metabolic disease.^8,9^ To accelerate the identification of individuals and disease states that would benefit from the future use of SARM1 inhibitors, we aimed to develop a minimally invasive screening method of SARM1 activity that can be applied in clinical studies.

Unlike neurons, peripheral blood mononuclear cells (PBMCs) represent an easily accessible population of cells for clinical screening. Prior research provides evidence of *SARM1* gene expression in hematopoietic lineage cells such as macrophages and T cells in mice, and monocytes, lymphocytes, natural killer cells, and T cells in humans.^10–16^ *SARM1* has also been detected in BlaER1, THP-1, HEK293T, SH-SY5Y, and NK-cell lines.^14,16^ However, the potential for quantification and tracking of SARM1 enzymatic activity in primary human PBMCs remains unknown.

We hypothesized that by pairing activators and inhibitors of SARM1 with analysis of downstream changes in cellular metabolites, we could quantify both the basal SARM1 activity and the SARM1 activation potential of human PBMCs. Activation of the SARM1 NADase occurs in response to injury, inflammation, and oxidative stress due to a high ratio of nicotinamide mononucleotide (NMN) to nicotinamide adenine dinucleotide (NAD+) within the cell and binding of NMN to the allosteric activation site of SARM1.^17^ This can be bypassed through direct SARM1 activation using either 3-acetylpyridine (3-AP) or pyrinuron, also known as Vacor, which are converted to 3-AP-MN and Vacor-MN by nicotinamide phosphoribosyltransferase (NAMPT) and bind SARM1 in lieu of NMN.^18,19^ The enzymatic cleavage of NAD^+^ through activated SARM1 subsequently creates byproducts including nicotinamide (NAM), cyclic ADP-ribose (cADPR), and ADP-ribose (ADPR), with previous work in neurons showing that cADPR accumulation is a sensitive and specific readout of SARM1 activity.^20^ To test this in PBMCs, we performed functional assays of SARM1 using known inhibitors and activators of its enzymatic activity.^18,19,21^ Specifically, human PBMCs were stimulated with SARM1 agonists 3-AP (1 mM) and Vacor (10, 100, 500 μM), for 4-hours to activate the enzymatic activity of SARM1. Agonist responses were compared relative to treatment with vehicle control for each independent participant. To confirm specificity for SARM1, a paired sample of cells within each group was simultaneously treated with 100 μM of SARM1 inhibitor DSRM-3716, which binds to the orthosteric site of SARM1 to inhibit its NADase activity.^21^ Our results reveal that Vacor drives a dose-dependent increase in cAPDR that is arrested when paired with SARM1 inhibitor DSRM-3716. Various changes in secondary metabolites were also characterized and reported herein, leading to a proposed working model for SARM1 function within PBMCs as it relates to NAD+ metabolism. Overall, these findings reveal that human PBMCs have detectable SARM1 activation potential and could be leveraged as a clinical readout of SARM1 expression and activity across diverse disease contexts.

## METHODS

### Human PBMCs

This study included PBMCs isolated from adolescent girls aged 12 to 16 years (n=7, mean 15.1±1.8 years). Informed consent and assent were obtained from the guardian and participants, respectively, under IRB #201908120. Participants were recruited from Washington University. Exclusion criteria included medical conditions including anemia, osteogenesis imperfecta, fetal alcohol syndrome, leukemia, congenital heart defects, neurological disorders, untreated hyper-or hypothyroidism, rheumatoid arthritis, hyperparathyroidism, epilepsy, Parkinson’s disease, cancer, coronary artery disease, peripheral vascular insufficiency, stroke, or premature ovarian failure. In addition, subjects using medications known to impact bone metabolism (bisphosphonates, glucocorticoids, GH) were excluded.

### RNA extraction and qPCR

Washed cells were resuspended in 600 μL of lysis buffer with freshly added 1% β-mercaptoethanol (β-ME) and homogenized with a 27-gauge needle. RNA was isolated using the Invitrogen PureLink RNA column (#12183018A) and eluted in 30 μL of nuclease-free water. RNA samples were stored at -80°C prior to analysis. RNA samples were transcribed to cDNA using the Maxima™ H Minus cDNA Synthesis kit (ThermoFisher Scientific, M1682) according to the manufacturer’s instructions. TaqMan quantitative PCR was used to measure gene expression levels in the cDNA. TaqMan™ Fast Advanced 2X Master Mix for qPCR (Applied Biosystems, # 4444557). *SARM1* expression was measured using the Hs00248344_m1 primer assay. The expression level was calculated based on a cDNA standard curve and compared to gene expression of ACTB (Hs99999903_m1) and HPRT1 (Hs02800695_m1).

### PBMC purification and treatment

Whole blood was collected from volunteer participants into three 6 mL K2 EDTA-treated vacutainers. The tubes were inverted multiple times to ensure complete mixing of the anticoagulant with the blood. Vacutainers were transported on ice and processed within 1 hour of collection. Approximately 15 mL of anticoagulant-treated blood was transferred to a 50 mL conical tube and diluted with 15 mL of room-temperature Dulbecco’s Phosphate Buffered Saline (DPBS). Separately, Ficoll-Paque media was mixed thoroughly by inverting the container. A 15 mL aliquot of the Ficoll-Paque media was withdrawn using a syringe and added to a new 50 mL conical tube. The diluted blood (30 mL) was gently layered onto the Ficoll-Paque media without mixing. The conical tube was centrifuged at 400 × g for 30 minutes at 18°C with no brake applied. This process resulted in distinct layers, including the plasma layer on top, the mononuclear cell layer in the middle, and Ficoll-Paque and red blood cells at the bottom. The plasma layer was carefully aspirated till the mononuclear cell layer. The mononuclear cells were then collected using a sterile Pasteur pipette, ensuring minimal contamination with Ficoll-Paque media or the remaining plasma. The collected mononuclear cells were transferred to a new 50 mL conical tube and washed with 30 mL of DPBS. The tube was centrifuged at 150 × g for 10 minutes at 18°C. After centrifugation, the supernatant was removed, and the cell pellet was resuspended in complete RPMI 1640 media supplemented with 10% patient-derived serum to achieve a final density of approximately 2 × 10^6^ cells/mL. Cells were divided into tubes for RNA isolation and metabolite extraction based on the number of treatment conditions. For example, if there were five treatment groups (Basal Condition, Vehicle, 10 μM DSRM-3716, 1 mM 3-AP, and 10 μM DSRM-3716 + 1 mM 3-AP), the cells were divided into ten 1.5 mL tubes, each containing approximately 2 million cells. The number of tubes and cell volumes were adjusted according to the total yield from the PBMC isolation process.

### Metabolite extraction

Washed cells were lysed in 160 μL ice cold 50% MS grade MeOH in H_2_O which was mixed thoroughly and incubated on ice for 5 minutes to extract metabolites. The 1.5 mL microcentrifuge tubes were centrifuged at 2,000 × g at 4°C for 5 minutes to pellet debris. The supernatant containing extracted metabolites was transferred to a new tube and mixed vigorously with 50 μL of HPLC-grade chloroform. The tubes were centrifuged at 20,000 × g at 4°C for 15 minutes. The clear upper aqueous phase was collected and stored at -80°C. Prior to LC-MS analysis, the samples were thawed and lyophilized under vacuum.

### Metabolite measurement using LC-MS/MS

Lyophilized samples were reconstituted with 70 μl of 5 mM ammonium formate and centrifuged at 12,000 x g for 10 min. Cleared supernatant were transferred to sample vials. Serial dilutions of standards for each metabolite in 5 mM ammonium formate were used for calibration. HPLC-mass spectrometry analysis was performed on an Agilent 1290 Infinity II liquid chromatography system (Agilent Technologies, Santa Clara, CA) with a flexible pump, multisampler, sample cooler and an MCT containing an Atlantis T3 column (2.1 × 150 mm, 3 μm) and VanGuard guard cartridge (2.1 mm X 5 mm, 3 μm) (Waters, Milford, MA), coupled to an Agilent 6470 Triple Quad mass spectrometer (Agilent Technologies, Santa Clara, CA). The mobile phase (0.15 ml/min) was 5 mM ammonium formate in water (A) and 100% methanol (B). The column was equilibrated with 0% B, maintained after injection for 2 min, then a linear gradient to 20% B applied over 4 min. The column was then ramped to 50% B over 2 min, and held at 50% for 2 min, then reverted back to 0% B over the next 5 min and allowed to re-equilibrate at 0% B for 9 min. The total run time was 24 min per sample. The injection volume was 10 μl. The mass spectrometer was equipped with an electrospray ion source which was operated in positive ion multiple reaction monitoring (MRM) mode for the detection of all metabolites. The [M+H]+ transitions were optimized for each metabolite and were selected as follows: m/z 560 → 136 for ADPR, m/z 348 → 136 for AMP, m/z 508 → 136 for ATP, m/z 542 → 428 for cADPR, m/z 349 → 137 for IMP, m/z 269 → 137 for Inosine, m/z 124 → 78 for NA, m/z 665 → 428 for NaAD, m/z 664 → 428 for NAD, m/z 666 → 649 for NADH, m/z 123 → 80 for Nam, m/z 336 → 124 for NaMN, m/z 335 → 123 for NMN, m/z 384 → 252 for s-AD and m/z 205 → 188 for TRP. The mass spectrometer settings for the fragmentation, the collision energy (CE) and the cell accelerator voltage were optimized for each of these transitions. Raw data were acquired and quantified using MassHunter Workstation software version B.08.00 for 6400 Series Triple Quadrupole (Agilent Technologies, Santa Clara, CA).

### Statistics

Biostatistical comparisons were performed in GraphPad Prism software. Differences in treatment-dependent changes between groups were evaluated by 2-way ANOVA with repeated measures for linked samples from each participant (e.g. treatment x SARM1 inhibitor). Post-hoc comparisons for column (agonist) and row (antagonist) effects were performed using Šídák’s multiple comparisons test. A P value less than 0.05 was considered significant with a P<0.10 interpreted as a trending result. For 2-way ANOVA, if no significant interaction term, significant individual effects of independent variables are presented; if the interaction was significant, this is presented in the figures.

## RESULTS

### *SARM1* is expressed in human PBMCs

*SARM1* gene expression in human PBMCs was measured with TaqMan qPCR. Amplification of *SARM1* based on the computed value threshold (C_T_) occurred after 31.7±0.2 cycles with linear decreases in amplification observed during serial dilution of a standard curve sample (Fig.1). For comparison, C_T_ values for housekeeping genes *HRPT1* and *ACTB* were 28.4±0.3 and 21.2±0.5, respectively with linear changes in amplification observed during serial dilution (Fig.1). This reaffirms positive *SARM1* expression in PMBCs, though at a lower level than standard housekeeping genes *HRPT1* and *ACTB*.

**Figure 1.**
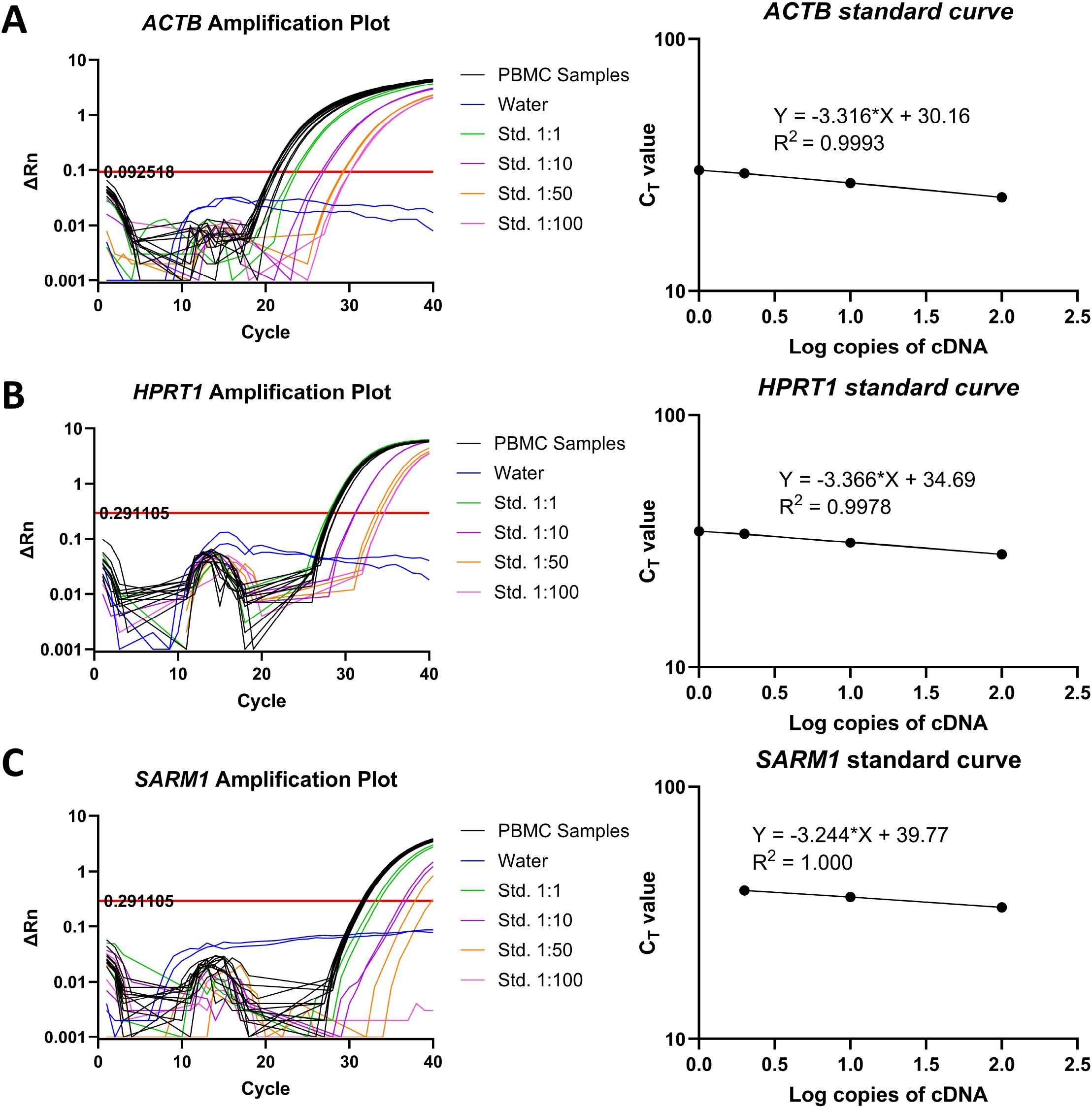
TaqMan qPCR amplification plots and standard curves of *SARM1* gene expression in human PBMCs. Gene expression was measured using TaqMan qPCR of human PBMC cDNA as well as a mix of cDNA for the standard curve. Amplification plots and standard curve plots are given for **(A)** *ACTB* expression, **(B)** *HPRT1* expression, and **(C)** *SARM1* expression. All samples and standard curves were run in duplicate and with n=6 PBMC samples. The left-side graphs have x-axis denoting the C_T_ cycle and the y-axis denoting change in normalized reporter (ΔRn) from the baseline signal. The Rn is calculated as the ratio of experimental fluorescence compared to the passive dye fluorescence (ROX), and the threshold (indicated by the red horizontal line) is determined as the level of fluorescence above background fluorescence. Right-side graphs are log regressions of standard curve C_T_ outputs versus log cDNA concentrations.

### cADPR is a functional readout of SARM1 activity in PBMCs

Activation of SARM1 increases the cellular pool of cADPR (Fig.2A).^20,22^ Consistent with this, cADPR was significantly elevated in PBMCs at even low concentrations of the SARM1 agonist Vacor (Fig.2B). Co-administration with SARM1 inhibitor DSRM-3716 prevented this increase, suggesting SARM1 dependence (Fig.2B). Unlike Vacor, SARM1 agonist 3-AP had minimal effects on cADPR in PBMCs. The related byproduct ADP-ribose (ADPR) was also regulated by Vacor in a SARM1 dependent manner, with noted suppression after 4-hours (Fig.2C).

**Figure 2.**
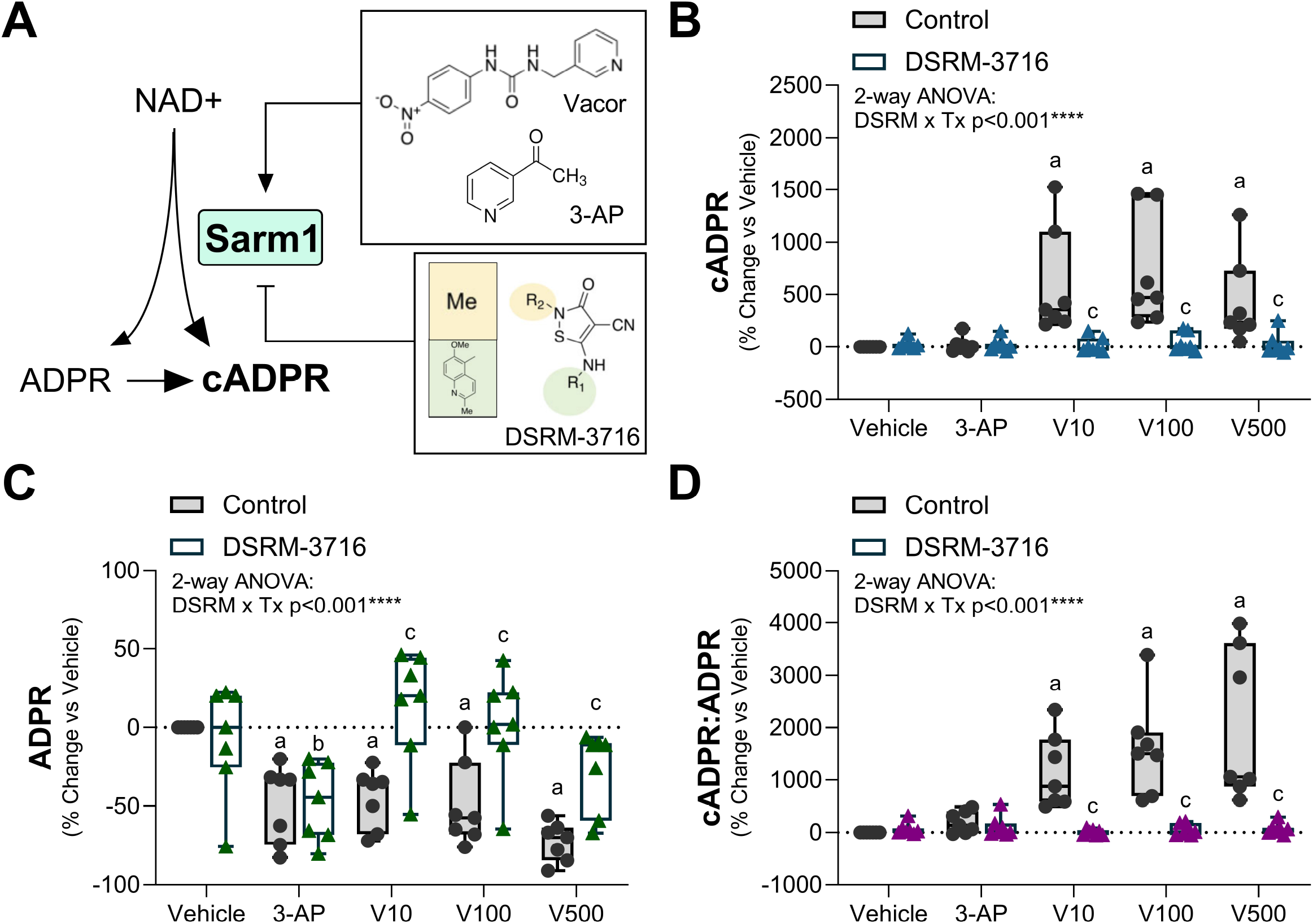
SARM1-dependent elevation of cADPR in human PBMCs. PBMCs from each participant were stimulated with SARM1 agonists 3-acetylpyridine (3-AP; 1 mM) and pyrinuron, also known as Vacor (V; 10, 100, 500 μM), for 4-hours to activate the enzymatic activity of SARM1. To test specificity for SARM1, a paired sample of cells within each group were simultaneously treated with 100 μM of SARM1 inhibitor DSRM-3716. Responses are expressed as a percent change vs each individual vehicle control. **(A)** Model of agonist and antagonist action on the SARM1 NADase enzyme. **(B)** Cyclic ADP-ribose (cADPR). **(C)** ADP-ribose (ADPR). **(D)** Ratio of cADPR:ADPR for each individual sample. 2-way ANOVA with repeated measures for linked samples from each participant with Šídák’s multiple comparisons test. ^a^p<0.05 vs Vehicle Control. ^b^ p<0.05 vs Vehicle DSRM-3716. ^c^p<0.05 vs Control within the same treatment group.

To consider flux from ADPR to cADPR, we also quantified the ratio of cADPR:ADPR in our samples (Fig.2D). The cADPR:ADPR ratio was substantially elevated with Vacor treatment and restored to baseline by co-administration of SARM1 inhibitor DSRM-3716. Minimal elevations were also noted with 3-AP, but this was not statistically significant. Overall, these findings show that SARM1 agonist Vacor potently and selectively stimulates the accumulation of cADPR. Our findings also suggest that utilization of the cADPR:ADPR ratio, in addition to cADPR alone, may hold value in tracking SARM1 activation in human PBMCs. Beyond this, it is important to note that treatment with DSRM-3716 alone in the vehicle control group did not significantly alter cADPR, ADPR, or the cADPR:ADPR ratio, suggesting minimal changes in these metabolites due to basal SARM1 activity in freshly isolated PMBCs (Fig.2B,C,D).

### SARM1 agonists deplete NAD+ and NMN with differential regulation of NaMN in PBMCs

When activated, the SARM1 NADase depletes cellular NAD+ and is regulated by metabolites upstream of NAD+ synthesis including NMN and nicotinic acid mononucleotide (NaMN). Physiologically, high NMN:NAD+ activates SARM1. This is bypassed by direct chemical SARM1 activators 3-AP and Vacor after their conversion to 3-AP-MN and Vacor-MN and binding to the allosteric site.^18,19^ NaMN can also competitively bind the allosteric site of SARM1 to block against activators NMN, 3-AP-MN, or Vacor-MN, inhibiting the activation of SARM1.^23^

Treatment with SARM1 activators 3-AP and Vacor significantly decreased NAD+ (Fig.3B). Partial restoration of NAD+ occurred with administration of SARM1 inhibitor DSRM-3716 in the Vacor groups only. This suggests the occurrence of SARM1-independent effects of both 3-AP and Vacor on NAD+ biosynthesis in PBMCs, as will be discussed below. SARM1 agonists 3-AP and Vacor also depleted NMN with partial restoration by DSRM-3716 in the 500 μM Vacor group only. SARM1 agonist 3-AP, but not Vacor, selectively elevated SARM1 inhibitor NaMN (Fig.3D). Elevated NaMN was also noted in the vehicle control group after treatment with SARM1 inhibitor DSRM-3716.

**Figure 3.**
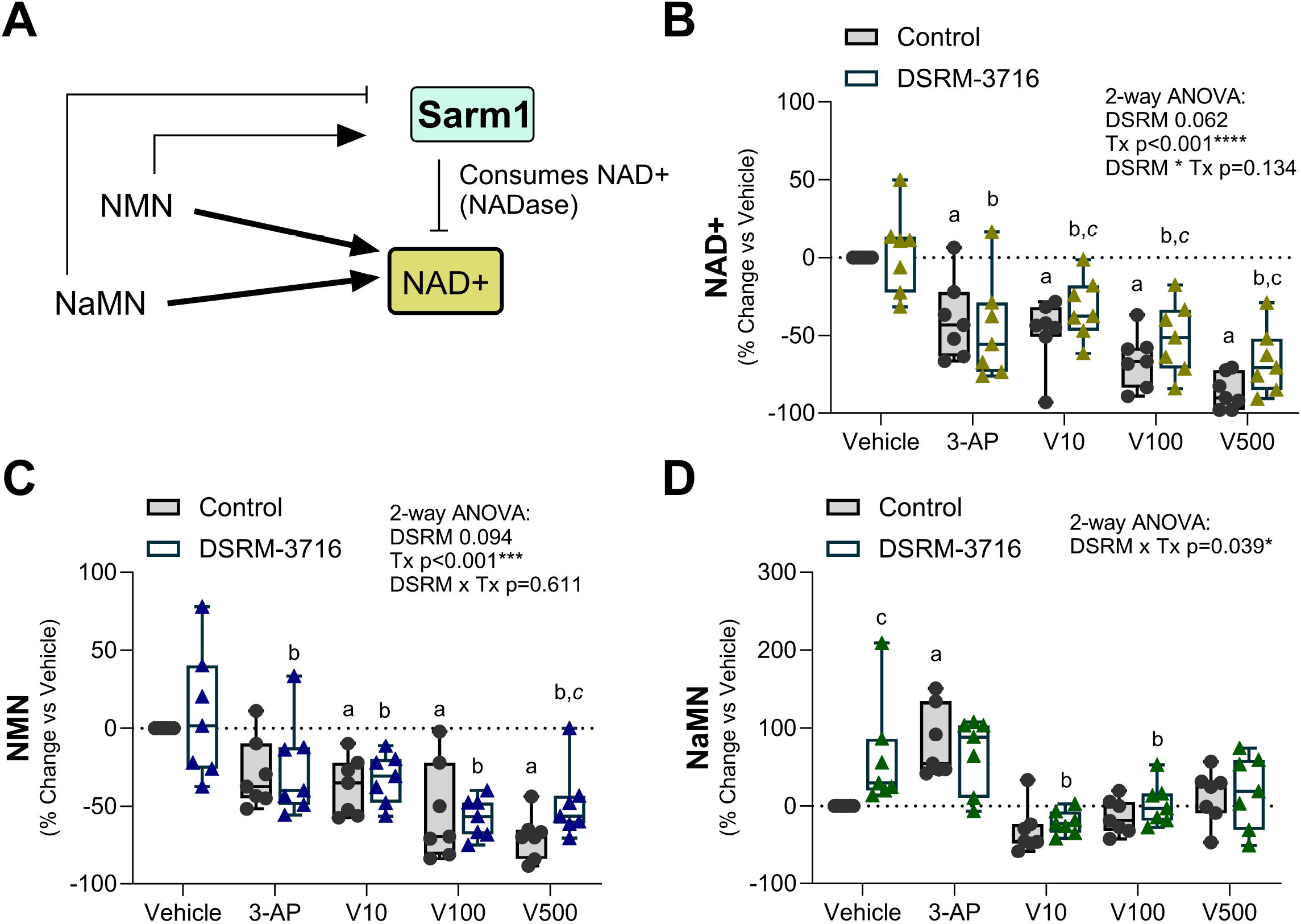
Generation and utilization of NAD+ after SARM1 activation in PBMCs. PBMCs from each participant were stimulated with SARM1 agonists 3-acetylpyridine (3-AP; 1 mM) and pyrinuron, also known as Vacor (V; 10, 100, 500 μM), for 4-hours to activate the enzymatic activity of SARM1. To test specificity for SARM1, a paired sample of cells within each group were simultaneously treated with 100 μM of SARM1 inhibitor DSRM-3716. Responses are expressed as a percent change vs each individual vehicle control. **(A)** Model of nicotinamide adenine dinucleotide (NAD+) production and consumption by the SARM1 NADase enzyme. **(B)** NAD+. **(C)** Nicotinamide mononucleotide (NMN). High NMN:NAD+ activates SARM1, but this is bypassed by 3-AP and Vacor which lead to direct activation of SARM1 independent of NMN. **(D)** Nicotinic acid mononucleotide (NaMN). High NaMN inhibits SARM1 and competes for binding of agonists 3-AP (as 3-AP-MN) and Vacor (as VacorMN). 2-way ANOVA with repeated measures for linked samples from each participant (n=7) with Šídák’s multiple comparisons test. ^a^p<0.05 vs Vehicle Control. ^b^p<0.05 vs Vehicle DSRM-3716. ^c^p<0.05 vs Control within the same treatment group. ^*a,b,c*^Same comparisons with p<0.1.

### Depletion of ATP and AMP occur at high doses of Vacor

We also examined changes in adenosine triphosphate (ATP) and adenosine monophosphate (AMP) by mass spec to track the health of the treated PBMCs (Fig.4A). Depletion of both ATP and AMP were observed in the high dose Vacor group only (500 μM), independent of SARM1 inhibition with DSRM-3716 (Fig.4B,C). Clumping and deterioration of PBMCs in this group was also noted upon microscopic examination (data not shown), suggesting that loss of ATP and AMP were likely secondary to drug toxicity and cell death.

**Figure 4.**
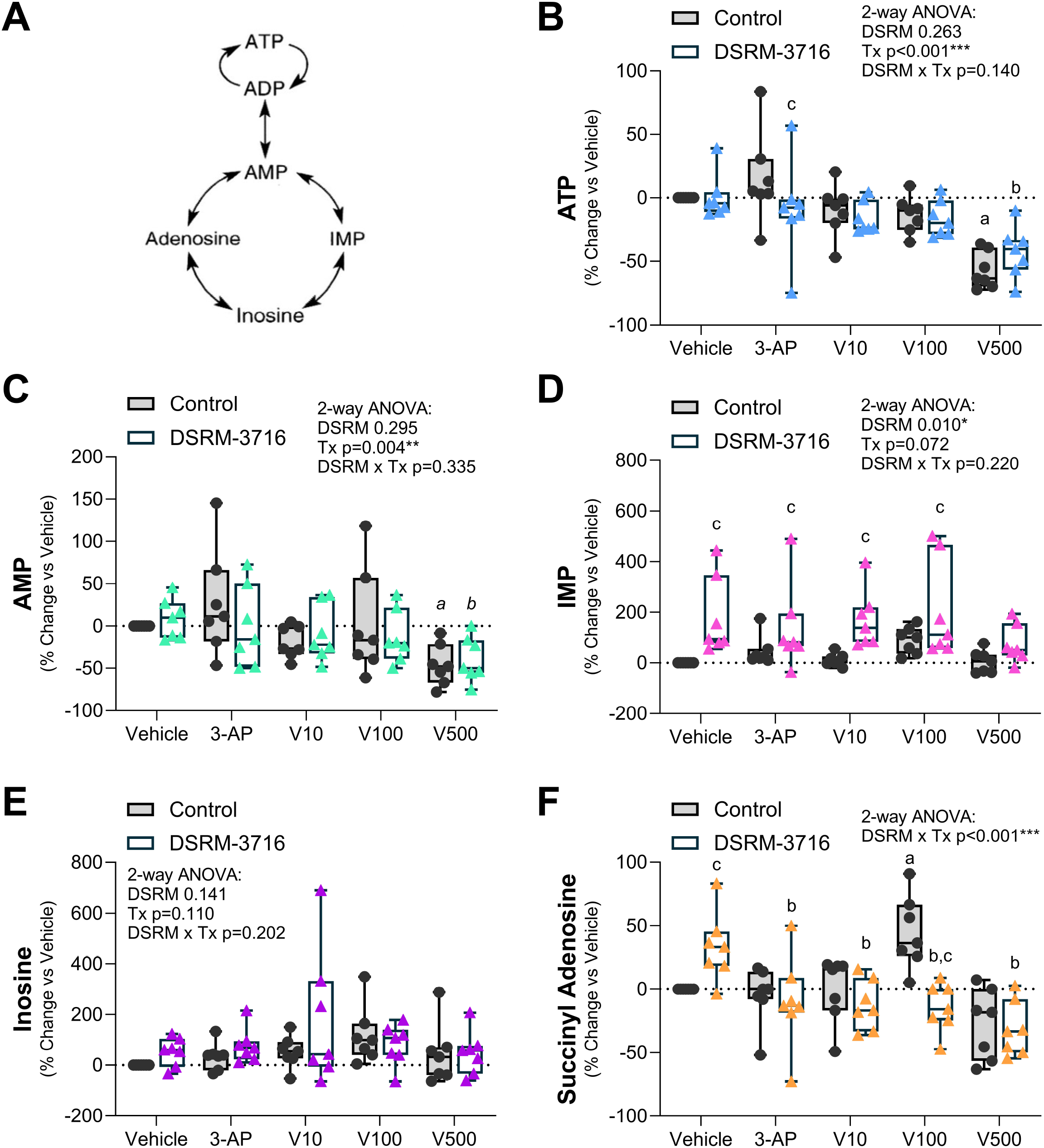
Tracking of ATP and downstream metabolites. PBMCs from each participant were stimulated with SARM1 agonists 3-acetylpyridine (3-AP; 1 mM) and pyrinuron, also known as Vacor (V; 10, 100, 500 μM), for 4-hours to activate the enzymatic activity of SARM1. To test specificity for SARM1, a paired sample of cells within each group were simultaneously treated with 100 μM of SARM1 inhibitor DSRM-3716. Responses are expressed as a percent change vs each individual vehicle control. **(A)** Model of ATP conversions and metabolic cycling. **(B)** ATP. **(C)** AMP. **(D)** IMP. **(E)** Inosine. **(F)** Succinyl adenosine. 2-way ANOVA with repeated measures for linked samples from each participant (n=7) with Šídák’s multiple comparisons test. ^a^p<0.05 vs Vehicle Control. ^b^p<0.05 vs Vehicle DSRM-3716. ^c^p<0.05 vs Control within the same treatment group. ^*a,b,c*^Same comparisons with p<0.1.

### Elevation of IMP occurs after treatment with SARM1 antagonist DSRM-3716

AMP is converted to metabolites including inosine monophosphate (IMP), inosine, and adenosine. Consistent elevation of the cellular IMP pool was observed after treatment with SARM1 inhibitor DSRM-3716 across all groups (Fig.4D). There were no changes in cellular inosine (Fig.4E) and variable shifts in succinyl adenosine in the vehicle and 100 μM Vacor groups only (Fig.4F).

## DISCUSSION

### Summary

Overall, our results indicate that human PBMCs have detectable expression of *SARM1* at the gene level. In addition, we observed functional SARM1 enzymatic activity and accumulation of cADPR in PBMCs after treatment with SARM1 agonist Vacor that was blocked by SARM1 inhibitor DSRM-3716. In neurons, cADPR has previously been shown to be a readout of SARM1 NADase activity with high sensitivity and specificity.^20^ This finding is of high interest because SARM1-dependent elevations in cADPR occur prior to the onset of irreversible cell death and may serve as an early marker of reversible SARM1 activity, possibly allowing for early detection and treatment intervention in the future. Of note, based on either cAPDR, ADPR, or the cADPR:ADPR ratio, SARM1 had limited basal activity in isolated PBMCs. This is similar to neurons which have very low basal activity, and suggests that clinical studies likely need to utilize both activators and inhibitors of SARM1 to reliably track SARM1 activation potential across different disease contexts.^20^

### Working model of SARM1 activation in PBMCs

Altogether, these data inform a working model of SARM1 activation in PBMCs (Fig.5) that is slightly different than what has been reported for neurons as it relates to changes in NAD+ and ADPR.^20^ First, the observation that SARM1 activators decrease NAD+ in manner that is only partially SARM1-dependent can be best explained if we consider (#1) the relatively low expression of *SARM1* in PBMCs and (#2) the direct regulation of upstream NAD+ biosynthesis through the known actions of 3-AP and Vacor on NAMPT and nicotinamide mononucleotide adenylyltransferase (NMNAT). In native form, 3-AP and Vacor compete with NAM as an alternative substrate for NAMPT.^18,19^ As observed in our data, this consequently decreases generation of NMN. Biosynthesis of NAD+ is then limited by both a reduction in precursor NMN and direct inhibition of NMNAT by Vacor-MN and 3-AP-MN. Vacor-MN and 3-AP-MN also increase NAD+ consumption by activating SARM1.^18,19^ Addition of orthostatic SARM1 inhibitor DSRM-3716 blocks NAD+ consumption by SARM1 but does not prevent the decrease in NAD+ biosynthesis, and we anticipate that this explains the only partial recovery of the PBMC NAD+ pool after treatment with orthostatic SARM1 inhibitor. Second, regarding the SARM1 agonist-dependent depletion of ADPR, we propose a model by which most cellular ADPR in PBMCs is generated by other highly expressed hematopoietic NADases including CD38 (Fig.5A). Previous work has shown that in HEK-293T cells, SARM1 produces cADPR from NADase activity at a more efficient rate than CD38 when activated, while CD38 NADase more often results in an ADPR byproduct.^24^ In our own data as well, activation of SARM1 in PBMCs appears to preferentially convert NAD+ to cADPR, as reported previously in neurons.^20^ We hypothesize that depletion of the NAD+ pool and SARM1-dependent generation of cADPR limits the availability of substrate for alternative NADase-based generation of ADPR, resulting in the observed decrease (Fig.5B). Future work will be needed to clarify this point.

**Figure 5.**
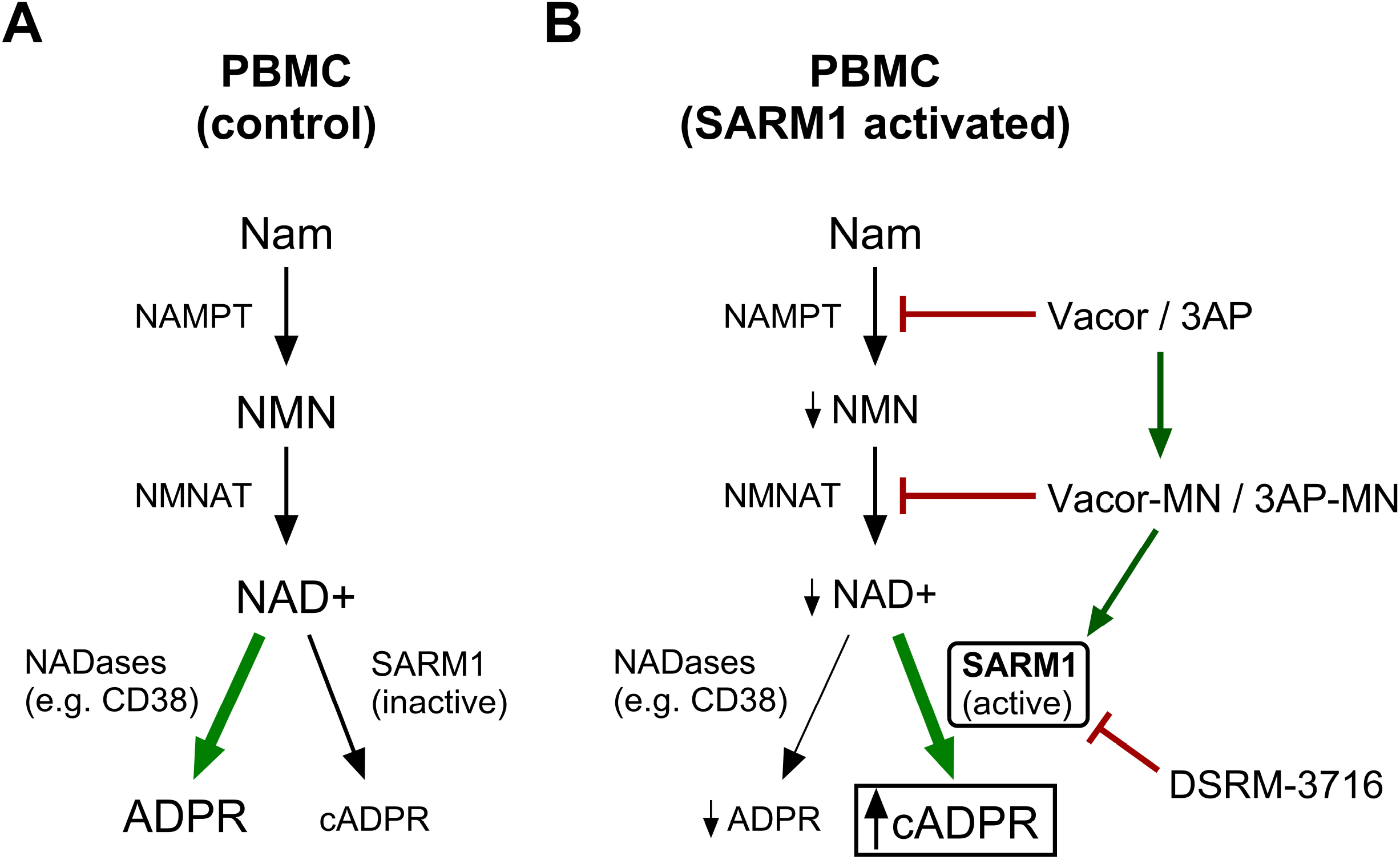
Working model of PBMC NAD+ metabolism and cADPR accumulation upon chemically induced SARM1 activation. Model for NAD+ metabolism in **(A)** control and **(B)** SARM1 activated PBMCs. The observation that SARM1 activators decrease NAD+ in manner that is only partially SARM1-dependent can be best explained if we consider (#1) the relatively low expression of *SARM1* in PBMCs relative to other NADases such as CD38 and (#2) the direct regulation of upstream NAD+ biosynthesis through the known actions of 3-AP and Vacor on NAMPT and NMNAT. In native form, 3-AP and Vacor compete with NAM as an alternative substrate for NAMPT. As observed in our data, this consequently decreases generation of NMN. Biosynthesis of NAD+ is then limited by both a reduction in precursor NMN and direct inhibition of NMNAT by Vacor-MN and 3-AP-MN. Vacor-MN and 3-AP-MN also increase NAD+ consumption by activating SARM1. Addition of orthostatic SARM1 inhibitor DSRM-3716 blocks NAD+ consumption by SARM1 but does not prevent the decrease in NAD+ biosynthesis, and we anticipate that this explains the only partial recovery of the PBMC NAD+ pool. Second, regarding the SARM1 agonist-dependent depletion of ADPR, we propose a model by which most cellular ADPR in PBMCs is generated by other highly expressed hematopoietic NADases such as CD38. Activation of SARM1 in PBMCs appears to preferentially convert NAD+ to cADPR, as reported previously in neurons. We propose that depletion of the NAD+ pool and SARM1-dependent generation of cADPR limits the availability of substrate for alternative NADase-based generation of ADPR, resulting in the observed decrease.

### Functions of SARM1 in PBMCs

Though this study was not intended to elucidate the function of SARM1 in PBMCs, some discussion of current literature is warranted. Functionally, SARM1 in hematopoietic cells has been hypothesized to mediate an anti-inflammatory response. Specifically, SARM1 has been hypothesized to be a negative regulator of toll-like receptor (TLR)-mediated production of IL-1β and TLR4 dependent TNF secretion.^10,12,16^ However, data from mouse studies report varying results on the role of SARM1 as a negative regulator of TLR signaling which suggests that the role and extent of SARM1 in the immune response may be distinct in mouse and human.^16,25,26^ Additionally, there is evidence that *SARM1* expression varies across mouse and human tissue with more *SARM1* expression in human peripheral tissues such as the liver and kidney while expression in mice is localized mainly to the nervous system.^13,27^ There have also been data suggesting that SARM1 localization within the cell varies based on species and cell-type.^26,28^ These variances in SARM1 location on the cell and tissue level could alter its function between mouse and human. Lastly, though SARM1 functions as a cytoplasmic NADase, one study has surprisingly detected SARM1 as a putative circulating factor in human blood serum.^29^ The current evidence for this is limited, but warrants additional investigation. Overall, it will be important to clarify the function of SARM1 across diverse human cell types and extracellular fluids, including PBMCs. Our study reveals that PBMCs are an easily accessible cell population to leverage for tracking SARM1 expression and activation in humans.

### Limitations and Additional Considerations

We found that SARM1 agonist 3-AP, which works well in neurons, did not fully activate SARM1 in PBMCs. This conclusion is based on the lack of changes in metabolite cADPR, which were readily observed with Vacor. One possible explanation for this is that 3-AP selectively elevated the cellular pool of SARM1 inhibitor NaMN in PBMCs, which may have led to autoinhibition of the SARM1 enzyme. This may also be due to the low potency of 3-AP-MN relative to Vacor-MN in binding to SARM1.^18,19^ The second unexpected result was the consistent elevation of IMP by DSRM-3716 regardless of treatment group. It may be that IMP can serve a readout of SARM1 inhibition in PBMCs. However, the mechanisms driving these changes in NaMN and IMP in PBMCs remain unknown.

### Conclusions

Altogether, we propose an assay for measuring SARM1 activation potential in human PBMCs by treating with optimal doses of SARM1 activator (100 μM Vacor) and inhibitor (100 μM DSRM-3716). As in neurons, our data also reveal that both cADPR and the cADPR:ADPR ratio can be used as a primary redout for SARM1 activity in PBMCs. Beyond this, we suggest quantifying NAD+ to monitor SARM1 NADase activity, NMN and NaMN to track the accumulation of SARM1 modulators, ATP/AMP as a readout of cellular health, and IMP as a putative marker of SARM1 inhibition.

## ACKNOWLEDGMENTS

This work was funded by grants from the National Institutes of Health including U24-DK115255 (ELS). We are grateful to Dr. Jeffrey Milbrandt and Dr. Aaron DiAntonio for their insightful comments and edits on this manuscript.

## BIBLIOGRAPHY

1. Waller, T.J., and Collins, C.A. (2022). Multifaceted roles of SARM1 in axon degeneration and signaling. Front. Cell. Neurosci. 16, 958900. 10.3389/fncel.2022.958900.

2. Loring, H.S., and Thompson, P.R. (2020). Emergence of SARM1 as a Potential Therapeutic Target for Wallerian-type Diseases. Cell Chem. Biol. 27, 1–13. 10.1016/j.chembiol.2019.11.002.

3. Figley, M.D., and DiAntonio, A. (2020). The SARM1 axon degeneration pathway: control of the NAD+ metabolome regulates axon survival in health and disease. Curr. Opin. Neurobiol. 63, 59–66. 10.1016/j.conb.2020.02.012.

4. Osterloh, J.M., Yang, J., Rooney, T.M., Fox, A.N., Adalbert, R., Powell, E.H., Sheehan, A.E., Avery, M.A., Hackett, R., Logan, M.A., et al. (2012). dSarm/Sarm1 is required for activation of an injury-induced axon death pathway. Science 337, 481–484. 10.1126/science.1223899.

5. Gerdts, J., Summers, D.W., Sasaki, Y., DiAntonio, A., and Milbrandt, J. (2013). Sarm1mediated axon degeneration requires both SAM and TIR interactions. J. Neurosci. 33, 13569– 13580. 10.1523/JNEUROSCI.1197-13.2013.

6. Yang, J., Wu, Z., Renier, N., Simon, D.J., Uryu, K., Park, D.S., Greer, P.A., Tournier, C., Davis, R.J., and Tessier-Lavigne, M. (2015). Pathological axonal death through a MAPK cascade that triggers a local energy deficit. Cell 160, 161–176. 10.1016/j.cell.2014.11.053.

7. Gerdts, J., Brace, E.J., Sasaki, Y., DiAntonio, A., and Milbrandt, J. (2015). SARM1 activation triggers axon degeneration locally via NAD^+^ destruction. Science 348, 453–457. 10.1126/science.1258366.

8. Pan, Z.-G., and An, X.-S. (2018). SARM1 deletion restrains NAFLD induced by high fat diet (HFD) through reducing inflammation, oxidative stress and lipid accumulation. Biochem. Biophys. Res. Commun. 498, 416–423. 10.1016/j.bbrc.2018.02.115.

9. Brazill, J.M., Shen, I.R., Craft, C.S., Magee, K.L., Park, J.S., Lorenz, M., Strickland, A., Wee, N.K., Zhang, X., Beeve, A.T., et al. (2024). Sarm1 knockout prevents type 1 diabetic bone disease in females independent of neuropathy. JCI Insight 9.

10. Carty, M., Goodbody, R., Schröder, M., Stack, J., Moynagh, P.N., and Bowie, A.G. (2006). The human adaptor SARM negatively regulates adaptor protein TRIF-dependent Toll-like receptor signaling. Nat. Immunol. 7, 1074–1081. 10.1038/ni1382.

11. Shanahan, K.A., Davis, G.M., Doran, C.G., Sugisawa, R., Davey, G.P., and Bowie, A.G. (2024). SARM1 regulates NAD+-linked metabolism and select immune genes in macrophages. J. Biol. Chem. 300, 105620. 10.1016/j.jbc.2023.105620.

12. Thwaites, R.S., Unterberger, S., Chamberlain, G., Gray, H., Jordan, K., Davies, K.A., Harrison, N.A., and Sacre, S. (2021). Expression of sterile-α and armadillo motif containing protein (SARM) in rheumatoid arthritis monocytes correlates with TLR2-induced IL-1β and disease activity. Rheumatology (Oxford) 60, 5843–5853. 10.1093/rheumatology/keab162.

13. Kim, Y., Zhou, P., Qian, L., Chuang, J.-Z., Lee, J., Li, C., Iadecola, C., Nathan, C., and Ding, A. (2007). MyD88-5 links mitochondria, microtubules, and JNK3 in neurons and regulates neuronal survival. J. Exp. Med. 204, 2063–2074. 10.1084/jem.20070868.

14. Panneerselvam, P., Singh, L.P., Selvarajan, V., Chng, W.J., Ng, S.B., Tan, N.S., Ho, B., Chen, J., and Ding, J.L. (2013). T-cell death following immune activation is mediated by mitochondria-localized SARM. Cell Death Differ. 20, 478–489. 10.1038/cdd.2012.144.

15. Doran, C.G., Sugisawa, R., Carty, M., Roche, F., Fergus, C., Hokamp, K., Kelly, V.P., and Bowie, A.G. (2021). CRISPR/Cas9-mediated SARM1 knockout and epitope-tagged mice reveal that SARM1 does not regulate nuclear transcription, but is expressed in macrophages. J. Biol. Chem. 297, 101417. 10.1016/j.jbc.2021.101417.

16. Sugisawa, R., Shanahan, K.A., Davis, G.M., Davey, G.P., and Bowie, A.G. (2024). SARM1 regulates pro-inflammatory cytokine expression in human monocytes by NADase-dependent and -independent mechanisms. iScience 27, 109940. 10.1016/j.isci.2024.109940.

17. Figley, M.D., Gu, W., Nanson, J.D., Shi, Y., Sasaki, Y., Cunnea, K., Malde, A.K., Jia, X., Luo, Z., Saikot, F.K., et al. (2021). SARM1 is a metabolic sensor activated by an increased NMN/NAD+ ratio to trigger axon degeneration. Neuron 109, 1118-1136.e11. 10.1016/j.neuron.2021.02.009.

18. Wu, T., Zhu, J., Strickland, A., Ko, K.W., Sasaki, Y., Dingwall, C.B., Yamada, Y., Figley, M.D., Mao, X., Neiner, A., et al. (2021). Neurotoxins subvert the allosteric activation mechanism of SARM1 to induce neuronal loss. Cell Rep. 37, 109872. 10.1016/j.celrep.2021.109872.

19. Loreto, A., Angeletti, C., Gu, W., Osborne, A., Nieuwenhuis, B., Gilley, J., Merlini, E., Arthur-Farraj, P., Amici, A., Luo, Z., et al. (2021). Neurotoxin-mediated potent activation of the axon degeneration regulator SARM1. eLife 10. 10.7554/eLife.72823.

20. Sasaki, Y., Engber, T.M., Hughes, R.O., Figley, M.D., Wu, T., Bosanac, T., Devraj, R., Milbrandt, J., Krauss, R., and DiAntonio, A. (2020). cADPR is a gene dosage-sensitive biomarker of SARM1 activity in healthy, compromised, and degenerating axons. Exp. Neurol. 329, 113252. 10.1016/j.expneurol.2020.113252.

21. Hughes, R.O., Bosanac, T., Mao, X., Engber, T.M., DiAntonio, A., Milbrandt, J., Devraj, R., and Krauss, R. (2021). Small Molecule SARM1 Inhibitors Recapitulate the SARM1-/-Phenotype and Allow Recovery of a Metastable Pool of Axons Fated to Degenerate. Cell Rep. 34, 108588. 10.1016/j.celrep.2020.108588.

22. Essuman, K., Summers, D.W., Sasaki, Y., Mao, X., DiAntonio, A., and Milbrandt, J. (2017). The SARM1 Toll/Interleukin-1 Receptor Domain Possesses Intrinsic NAD+ Cleavage Activity that Promotes Pathological Axonal Degeneration. Neuron 93, 1334-1343.e5. 10.1016/j.neuron.2017.02.022.

23. Sasaki, Y., Zhu, J., Shi, Y., Gu, W., Kobe, B., Ve, T., DiAntonio, A., and Milbrandt, J. (2021). Nicotinic acid mononucleotide is an allosteric SARM1 inhibitor promoting axonal protection. Exp. Neurol. 345, 113842. 10.1016/j.expneurol.2021.113842.

24. Zhao, Z.Y., Xie, X.J., Li, W.H., Liu, J., Chen, Z., Zhang, B., Li, T., Li, S.L., Lu, J.G., Zhang, L., et al. (2019). A Cell-Permeant Mimetic of NMN Activates SARM1 to Produce Cyclic ADP-Ribose and Induce Non-apoptotic Cell Death. iScience 15, 452–466. 10.1016/j.isci.2019.05.001.

25. Carty, M., Kearney, J., Shanahan, K.A., Hams, E., Sugisawa, R., Connolly, D., Doran, C.G., Muñoz-Wolf, N., Gürtler, C., Fitzgerald, K.A., et al. (2019). Cell Survival and Cytokine Release after Inflammasome Activation Is Regulated by the Toll-IL-1R Protein SARM. Immunity 50, 1412-1424.e6. 10.1016/j.immuni.2019.04.005.

26. Carty, M., and Bowie, A.G. (2019). SARM: From immune regulator to cell executioner. Biochem. Pharmacol. 161, 52–62. 10.1016/j.bcp.2019.01.005.

27. Mink, M., Fogelgren, B., Olszewski, K., Maroy, P., and Csiszar, K. (2001). A novel human gene (SARM) at chromosome 17q11 encodes a protein with a SAM motif and structural similarity to Armadillo/beta-catenin that is conserved in mouse, Drosophila, and Caenorhabditis elegans. Genomics 74, 234–244. 10.1006/geno.2001.6548.

28. Peng, J., Yuan, Q., Lin, B., Panneerselvam, P., Wang, X., Luan, X.L., Lim, S.K., Leung, B.P., Ho, B., and Ding, J.L. (2010). SARM inhibits both TRIF- and MyD88-mediated AP-1 activation. Eur. J. Immunol. 40, 1738–1747. 10.1002/eji.200940034.

29. Alrawaili, M.S., Abuzinadah, A.R., AlShareef, A.A., Hindi, E.A., Bamaga, A.K., Alshora, W., and Sindi, H. (2024). Serum SARM1 Levels and Diabetic Peripheral Neuropathy in Type 2 Diabetes: Correlation with Clinical Neuropathy Scales and Nerve Conduction Studies and Impact of COVID-19 vaccination. Vaccines (Basel) 12. 10.3390/vaccines12020209.

